# Acidosis Licenses the NLRP3 Inflammasome-Inhibiting Effects of Beta-Hydroxybutyrate and Short-Chain Carboxylic Acids

**DOI:** 10.1101/2025.05.01.650510

**Authors:** Madeleine M. Mank, Kaitlyn A. Zoller, V. Amanda Fastiggi, Jennifer L. Ather, Matthew E. Poynter

**Affiliations:** Department of Medicine, Division of Pulmonary Disease and Critical Care, University of Vermont, and The Vermont Lung Center, Burlington, VT, 05405, USA; Cellular, Molecular, and Biomedical Sciences Doctoral Program, University of Vermont, Burlington, VT, 05405, USA

**Author notes:** Correspondence: Matthew E. Poynter, PhD, Department of Medicine, Pulmonary Disease and Critical Care, University of Vermont, Firestone Medical Research Building 262, 149 Beaumont Ave, Burlington, VT 05405. MMM and MEP designed research studies, conducted experiments, acquired data, analyzed results, prepared figures, and wrote the manuscript. KAZ, VAF, JLA, and MEP conducted experiments, acquired data, and analyzed results. All authors edited the manuscript and approved the final version. This work was funded by US National Institute of Health research grants R01 HL142081, R21 AI186022, and T32 HL076122, research grants from the University of Vermont (UVM) Larner College of Medicine and Department of Medicine, research grants from the UVM Office of Undergraduate Research, and fellowship grants from the US Department of Education (the CMB-Graduate Assistance in Areas of National Need (CMB-GAANN) Training Grant and the Vermont Space Grant Consortium (VTSGC) Graduate Fellowship Program.

**Keywords:** Beta-hydroxybutyrate, Inflammation, Ketone Bodies, Macrophage, NLRP3 inflammasome

## Abstract

NLRP3 inflammasome activation induces the cleavage and secretion of IL-1β and IL-18, and causes pyroptosis. Generated during times of energetic crisis (*e.g.*, caloric insufficiency), the ketone body β-hydroxybutyrate (BHB) has been reported to inhibit NLRP3 inflammasome activation. However, the conditions under which BHB exerts this activity and whether other short-chain carboxylic acids (SCCAs) share this effect are unexplored. Since BHB is often produced in high abundance endogenously accompanied by metabolic acidosis, we aimed to examine the pH-dependence for the ability of BHB and similar molecules to inhibit NLRP3 inflammasome activation and to test receptors conferring these effects. Whereas β-hydroxybutyric acid (BHBA) enantiomers function equivalently to dose-dependently inhibit NLRP3 inflammasome-induced IL-1β secretion, sodium-β-hydroxybutyrate (NaBHB) and NaOH-neutralized BHBA do not inhibit NLRP3 inflammasome activation. Acidifying the pH of the NaBHB stock solution or the media in which cells are exposed to NaBHB, or allowing the cells to endogenously acidify their media, enables NaBHB to inhibit NLRP3 inflammasome activation. Several other SCCAs also inhibit NLRP3 inflammasome activation in a pH-dependent manner and prevent pyroptotic cell death. Finally, Free Fatty Acid Receptor 3 (GPR41/FFAR3) activation phenocopies and augments the NLRP3 inflammasome-inhibiting effects of BHBA. In conclusion, acidification licenses the ability of BHB and related SCCAs to inhibit NLRP3 inflammasome activation, in part through GPR41/FFAR3, thereby expanding the repertoire of metabolites capable of modulating this important pro-inflammatory pathway during times of energetic crisis and optimizing conditions for the potential use of ketone bodies as anti-inflammatories.

**Graphical Abstract:** 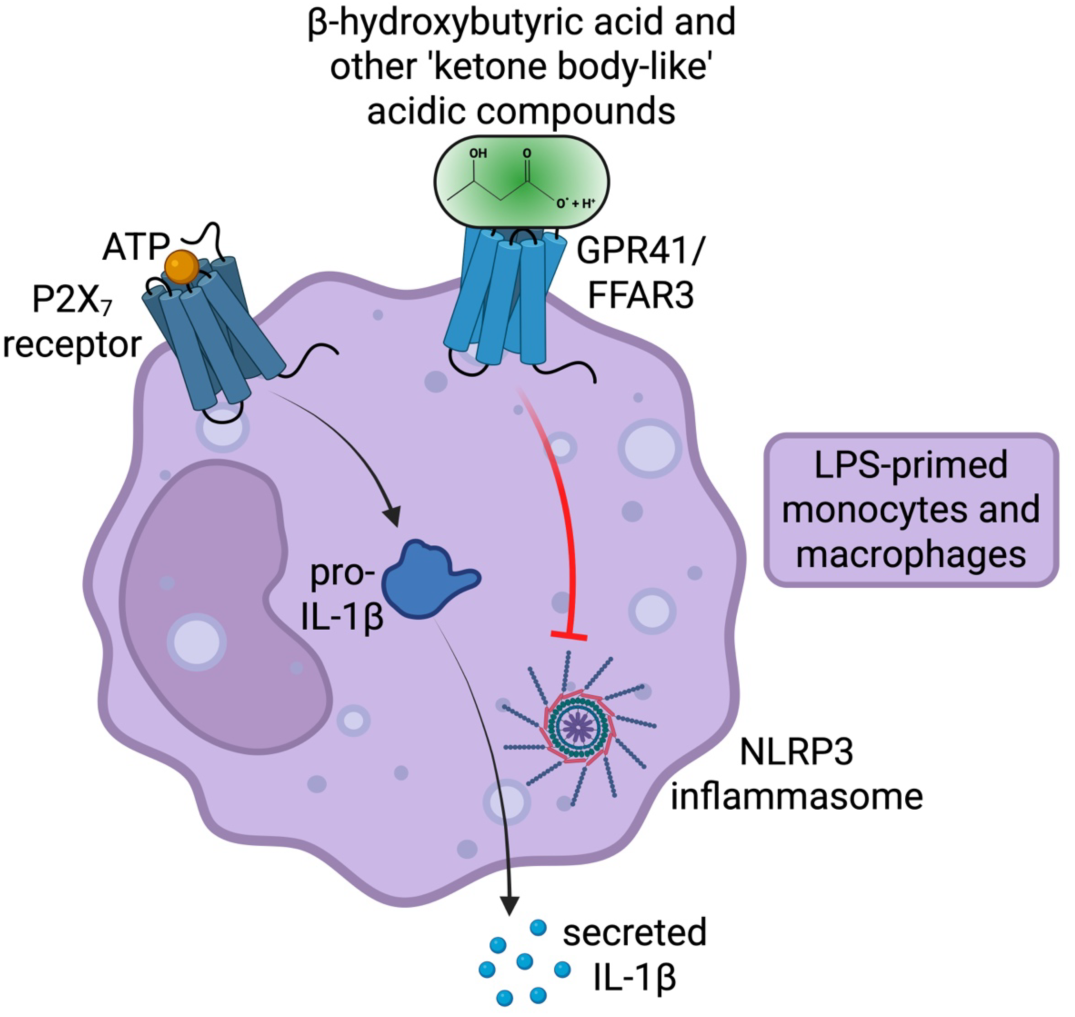

## Introduction

Inflammasomes are a family of large intracellular multi-protein pattern recognition receptor (PRR) complexes that respond to a wide variety of exogenous pathogen associated molecular patterns (PAMPs) and endogenous danger associated molecular patterns (DAMPs) from the host or generated through homeostatic disturbances (1, 2). The activation and oligomerization of the inflammasome complex initiates caspase-1 activation, which proteolytically processes the pro-inflammatory cytokines, IL-1β and IL-18, facilitating their secretion, and cleaves the pore-forming effector gasdermin D (GSDMD), which induces a form of inflammatory cell death known as pyroptosis (1, 2). Stimulation of a functional inflammasome requires two steps: priming and activation. During priming, activation of the transcription factor NF-κB, typically a consequence of PRR-initiated signaling, leads to the production of several inflammasome components and secretion of the pro-inflammatory cytokine TNFα (3). Inflammasome activation requires the exposure of cells to a separate set of PAMPs and DAMPs, which initiate oligomerization of one of several different Nucleotide Oligomerization Domain (NOD)-Like Receptor (NLR) proteins, the adaptor protein Apoptosis-associated Speck-like protein containing a CARD (ASC), and pro-caspase-1 into an organized inflammasome complex (4). ASC is phosphorylated at specific tyrosine residues for its activation (5), promoting the formation of a discrete ASC speck within stimulated cells that functions as a platform for efficient pro-IL-1β and pro-IL-18 cleavage (6). These cytokines go on to promote vasodilation, attract and stimulate neutrophils, induce fever, and activate the acute phase response within an organism (7). The final outcome of inflammasome activation, GSDMD-mediated pyroptotic cell death, is understood to amplify the immune response while depleting pathogens of their host leukocyte niche (8).

Activation of the NLRP3 inflammasome can occur in response to a diverse set of PAMPs and DAMPs, such as ATP, nigericin, alum, asbestos, silica, and cholesterol crystals (9–13). These agonists activate the inflammasome through a multitude of different mechanisms, including membrane pore formation and lysosomal rupture, iron fluxes, reactive oxygen species (ROS) production, and post-translational modifications, which eventually converge on K^+^ efflux and ASC multimerization (5, 14–20). Expressed predominantly by macrophages, the NLRP3 inflammasome plays a major function in immune homeostasis (21). While the NLRP3 inflammasome confers protection against pathogens, its excessive activation is linked to the development of several diseases, such as atherosclerosis, diabetes, gout, and multiple sclerosis (22–24).

During illnesses ranging from mild to critical, metabolic disturbances such as lactic acidosis, altered mitochondrial function, and impaired nutrient utilization can significantly affect immune responses and systemic homeostasis. One adaptive response to energetic stress is the increased production of ketone bodies, which serves not only as an alternative energy substrate but also as a signaling molecule with immunomodulatory properties. Ketone bodies can become elevated systemically as fatty acids consumed in the diet (25) or mobilized from adipose tissue as a consequence of energetic demand (26–28) are catabolized through β-oxidation in the liver to acetoacetate (AcAc) and β-hydroxybutyrate (BHB), which are then released into the circulation and can be used as an energy source by cells throughout the body (29). BHB comprises approximately 78% of the circulating concentration of ketone bodies and can be elevated during energetic crises such as critical illness (30, 31) or during times of low carbohydrate availability, such as during fasting (32) or prolonged exercise (33). BHB can also be elevated through the consumption of particular foods; for example, high-fat, low-carbohydrate diets provide ketogenic substrates that lead to the production of BHB (25, 34, 35). Additionally, ketone body precursors such as 1,3-butanediol (1,3-BD) (36), or ketone esters, a dietary supplement approved for human use (37), can transiently elevate ketone body concentrations. Importantly, ketone body augmentation in human subjects is well-tolerated (34).

In ill patients, elevated BHB levels may reflect an adaptive metabolic shift aimed at preserving cellular function under energetic stress (30). Importantly, BHB exerts anti-inflammatory effects, including suppression of NF-κB activation (38) and inhibition of the NLRP3 inflammasome and subsequent IL-1β production (39–42), which occurs independently of its role as a metabolic fuel and is partly mediated through reduction of potassium efflux and ROS production. As an energy source, ketone bodies make cells less reliant on glycolysis (29, 35, 43–46) and therefore produce less lactic acid, a catabolite implicated as a causal factor in many diseases. Ketone bodies have been reported to function through cell surface receptors, including the G protein-coupled receptors hydroxycarboxylic acid receptor 2 (HCAR2/GPR109a) and free fatty acid receptor 3 (FFAR3/GPR41) (26, 27, 47–49).

We hypothesized that, considering endogenous ketone generation is oftentimes accompanied by acidosis, the inflammasome-modulating effects of BHB, itself a carboxylic acid, may be influenced by pH. Our objectives were to evaluate the involvement of pH on the NLRP3-inhibiting effects of BHB and other ketone body-like short-chain carboxylic acids, as well as to identify mechanisms through which they modulate these effects. Further understanding the efficacy and mechanisms of ketone bodies *in vitro* could provide new insight into their functions endogenously during energetic crises, such as in critical illness, and enable optimization of their potential for therapeutic approaches to control inflammatory conditions. Understanding the precise conditions under which BHB contributes to recovery or pathology may open new avenues for metabolic interventions in sepsis, trauma, and other critical care contexts, as well as in chronic diseases.

## Materials and Methods

### Study approval

Studies involving potentially hazardous materials were reviewed and approved by the University of Vermont’s Institutional Biosafety Committee (protocol REG201900052).

### Reagents

Ultra Pure *E. coli* O111:B4 LPS and ATP were purchased from Invivogen (San Diego, CA). (*R,S*)-β-hydroxybutyric acid, (*R*)-β-hydroxybutyric acid, (*S*)-β-hydroxybutyric acid, α-hydroxybutyric acid, butyric acid, sodium butyrate, 4-amino-3-hydroxybutyric acid, sodium propanoate, 2-hydroxypropanoic acid, 3-hydroxypropanoic acid, 3-hydroxyvaleric acid, and 2-hydroxyhexanoic acid were purchased from Sigma-Aldrich (St. Louis, MO). A separate source of (*R,S*)-β-hydroxybutyric acid was purchased from Alfa Aesar through Fisher Scientific (Hampton, NH). Hydrochloric acid (HCl), nicotinic acid, and sodium nicotinate were purchased from Sigma-Aldrich, sodium hydroxide (NaOH) was purchased from Thermo Fisher Scientific (Waltham, MA), DMSO was purchased from MP Biomedicals (Santa Ana, CA), digitonin was purchased from Promega Corporation (Madison, WI), AR420626 was purchased from Cayman Chemical Company (Ann Arbor, MI), and AZD3965 was purchased from Med Chem Express (Monmouth Junction, NJ).

### Mouse macrophages

J774.1 murine macrophages purchased from American Type Culture Collection (Manassas, VA) were cultured at 37°C in 95% humidified air containing 5% CO_2_ using DMEM (Thermo Fisher Scientific) containing 10% FBS, 2 mM L-glutamine, 50 U/ml penicillin, and 50 μg/ml streptomycin (Life Technologies, Grand Island, NY). Cells were not used beyond passage 20 to reduce variability between the experiments. For experiments, J774 cells were plated at 1×10^6^ cells/ml in 125 μl media in 96-well plates and allowed to grow overnight. The following day, the media were removed, 100 μl of fresh media were added, and cells were treated as indicated within the figure legends for each experiment using 50 ng/ml LPS (a dose previously determined to elicit half-maximal IL-1β secretion upon ATP-induced stimulation) and 5 mM ATP in the absence or presence of ketone bodies.

### Human PBMCs

Heparinized blood collected from normal volunteers who were enrolled in a study approved by the Institutional Review Board (protocol 18-0600) and who provided written informed consent was diluted 1:1 in PBS, and 4 ml of diluted blood was layered over 3ml of lymphocyte separation medium (LSM) (MP Biomedicals) then centrifuged at 400 x *g* for 15 minutes at room temperature. The top layer of plasma was aspirated and discarded, leaving 2-3mm above the buffy coat, and the buffy coat and half the lower LSM layer was aspirated, mixed with PBS and centrifuged at 200 x *g* for 10 minutes at room temperature. The resuspended cells were cryopreserved at 1×10^7^ cells/ml in Recovery Cell Freezing Medium (Thermo Fisher Scientific), slowly chilled to −80°C, and stored in the vapor phase of liquid nitrogen. Thawed cells were washed once in PBS, resuspended in DMEM containing 10% FBS, 50 U/ml penicillin, and 5 μg/ml streptomycin, and plated at 1×10^6^ cells/ml. After an overnight incubation, cells were treated with Ultra Pure *E. coli* O111:B4 lipopolysaccharide at 10 ng/mL (a dose determined in previous studies to elicit half-maximal TNFα production) for 4 h and incubated with ketone bodies for 10 min before being stimulated with 5mM ATP for 1 h to induce NLRP3 inflammasome activation and IL-1β secretion.

### Cytokine immunoassays

Conditioned media from cell culture studies were collected at the indicated timepoints, centrifuged at 3300 x *g* to pellet cellular debris, transferred into new tubes or multi-well plates, and frozen at −20°C until analysis. ELISAs to quantitate mouse TNFα and IL-1β, as well as human TNFα, IL-1β, and IL-18, were DuoSets from R&D Systems (Minneapolis, MN) used according to manufacturer-recommendations with samples diluted to coincide with the range of the standards. Optical densities were read using a Bio-Tek Instruments (Winooski, VT) Synergy HTX multimode plate reader at 450 nm with background subtraction at 570 nm.

### Cell health measurements

The pH of conditioned media from J774.1 cells was measured using a FiveEasy pH/mV bench meter with sensor (Mettler Toledo, Columbus, OH) immediately after removal from the CO_2_ incubator. Cell viability and cytotoxicity were measured by staining scraped cells with trypan blue and counting clear and blue cells by light microscopy on a hemocytometer or by using the CytoTox96 Non-Radioactive Cytotoxicity Assay, based on lactate dehydrogenase (LDH) activity, according to manufacturer’s protocols (Promega).

### Data acquisition, data availability, and statistical analysis

All experiments were conducted using 3-4 replicates per condition and were repeated at least three times. Representative results are presented. The datasets generated during and/or analyzed during the current study are available from the corresponding author upon reasonable request. Data were analyzed by Brown-Forsythe and Welch ANOVA tests followed by Dunnett’s T3 multiple comparisons tests, with individual variances computed for each comparison using GraphPad Prism 10.4.2 (534) for macOS (GraphPad Software, Inc., La Jolla, CA; RRID:SCR_002798). Data presented are mean ± SEM from a representative experiment. A p-value smaller than 0.05 in the multiple comparisons post-hoc test was considered statistically significant. The significance levels of the tested comparisons are indicated in the figure legends.

## Results

### NLRP3 inflammasome-stimulated IL-1β secretion from mouse macrophages is inhibited by the ketone body β-hydroxybutyric acid but not by its sodium salt, sodium β-hydroxybutyrate

Ketone bodies, particularly β-hydroxybutyrate (BHB), have been reported to exert anti-inflammatory effects, especially via inhibition of NLRP3 inflammasome activation (39–42). Using LPS-primed J774.1 mouse macrophages that we have reported to be susceptible to NLRP3 inflammasome inhibition by several short-chain alcohols (50) and by β-hydroxybutyric acid (BHBA) (51), we confirmed that when an enantiomeric mixture of BHBA ((*R,S*)-BHBA) or single BHBA enantiomers ((*R*)-BHBA or (*S*)-BHBA), or (*R,S*)-BHBA from another supplier, were added immediately preceding ATP-induced NLRP3 inflammasome activation TNFα concentrations were relatively unaffected whereas (**Figure 1A**) there was a dose-dependent decrease in the abundance of IL-1β that accumulated in the culture medium (**Figure 1B**). To assess the potential for BHBA to cause interference in the ELISAs, we supplemented standards and culture media with BHBA at concentrations up to 20 mM. This resulted in a modest (up to 20%) reduction in detected TNFα and a minor (5%) reduction in IL-1β, indicating that observed differences in TNFα likely reflect assay interference, whereas the reductions in IL-1β are attributable to BHBA’s biological activity. In contrast to BHBA, the mixed enantiomer sodium salt of BHB, sodium BHB (NaBHB) had no effect on the abundance of TNFα (**Figure 1C**) or IL-1β (**Figure 1D**) secreted from LPS+ATP-stimulated J774.1 macrophages.

**Figure 1.**
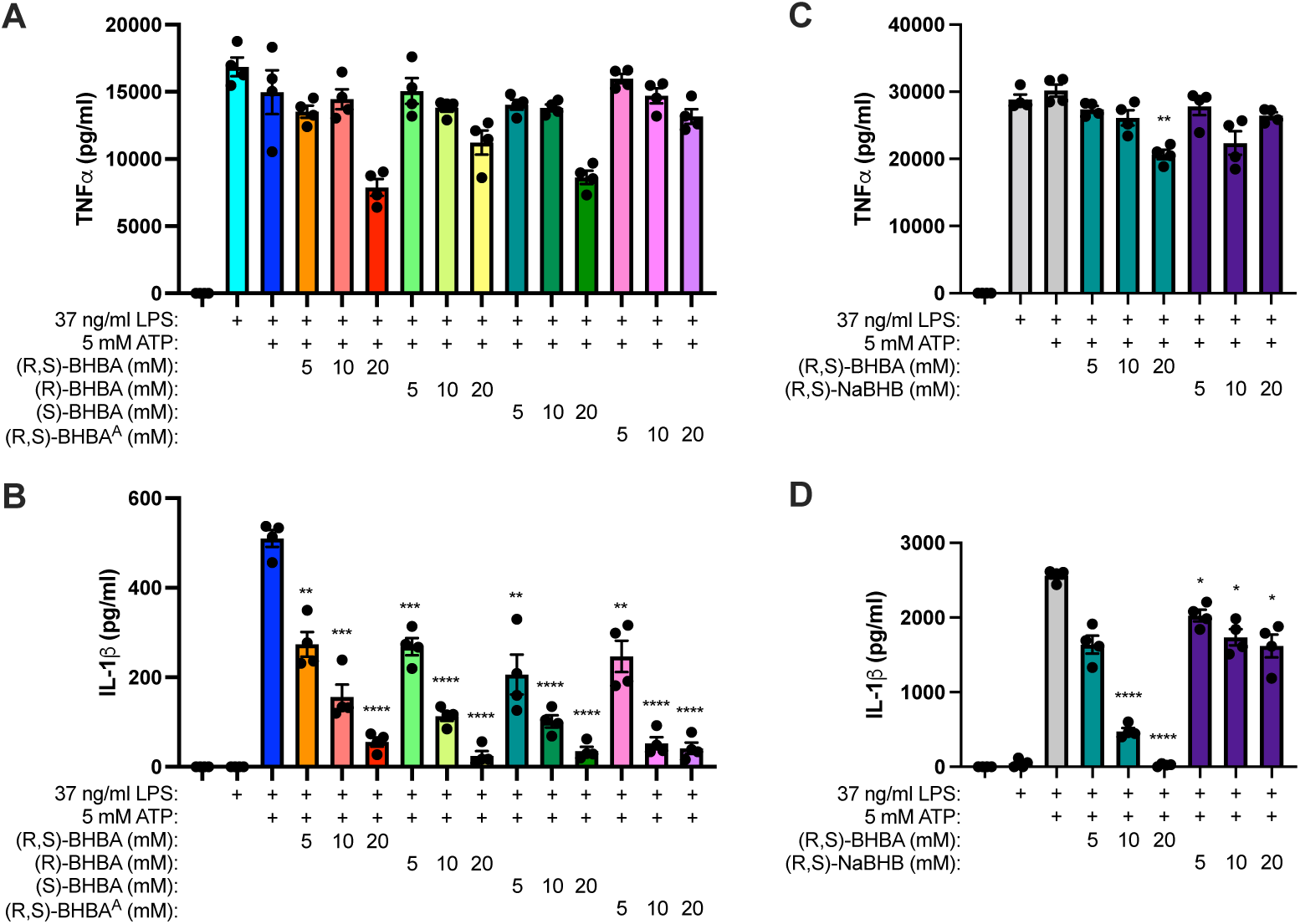
β-hydroxybutyric acid (BHBA) enantiomers function equivalently to inhibit NLRP3 inflammasome activation, whereas sodium-β-hydroxybutyrate (NaBHB) does not inhibit NLRP3 inflammasome activation. J774.1 macrophages were unstimulated or primed with LPS for 6 hours then stimulated with ATP in the presence of increasing concentrations of a racemic mixture (*R,S*)-of β-hydroxybutyric acid (BHBA), (*R*)-BHBA, or (*S*)-BHBA from Sigma-Aldrich, or a racemic mixture of BHBA from Alfa Aesar (BHBA^A^) for 1 hour, after which soluble TNFα **(A)** and IL-1β **(B)** were measured from the cell culture supernatants. J774.1 macrophages were unstimulated or primed with LPS for 6 hours then stimulated with ATP in the presence of increasing concentrations of (*R,S*)-BHBA, or (*R,S*)-sodium β-hydroxybutyrate (NaBHB) for 1 hour, after which soluble TNFα **(C)** and IL-1β **(D)** were measured from the cell culture supernatants. N = 4 samples/group. * = p≤0.05, ** = p≤0.01, *** = p≤0.001, **** = p≤0.0001 compared to LPS+ATP.

### Acidifying the pH of NaBHB confers upon it the capacity to inhibit NLRP3 inflammasome activation and acid alone is insufficient to decrease the abundance of soluble IL-1β

To determine the impact of pH on the capacity of BHB to inhibit NLRP3 inflammasome activation, a 1M stock of NaBHB was adjusted to more acidic pH using HCl and then added to LPS-primed J774.1 macrophages immediately before ATP-induced inflammasome activation. The abundance of TNFα as not substantially affected by BHBA, NaBHB, or acidified NaBHB (**Figure 2A**). In contrast, whereas BHBA strongly inhibited IL-1β release in a dose-dependent manner and NaBHB did not, acidified NaBHB displayed an increased capacity to inhibit IL-1β secretion as the pH was made increasingly acidic (pH6 < pH4 < pH2) (**Figure 2B**). Adding an equivalent amount of HCl alone was unable to substantially affect the abundance of TNFα (Figure 2C) or IL-1β detected, indicating that both an acidic environment and BHB are required for the NLRP3 inflammasome-inhibiting activity.

**Figure 2.**
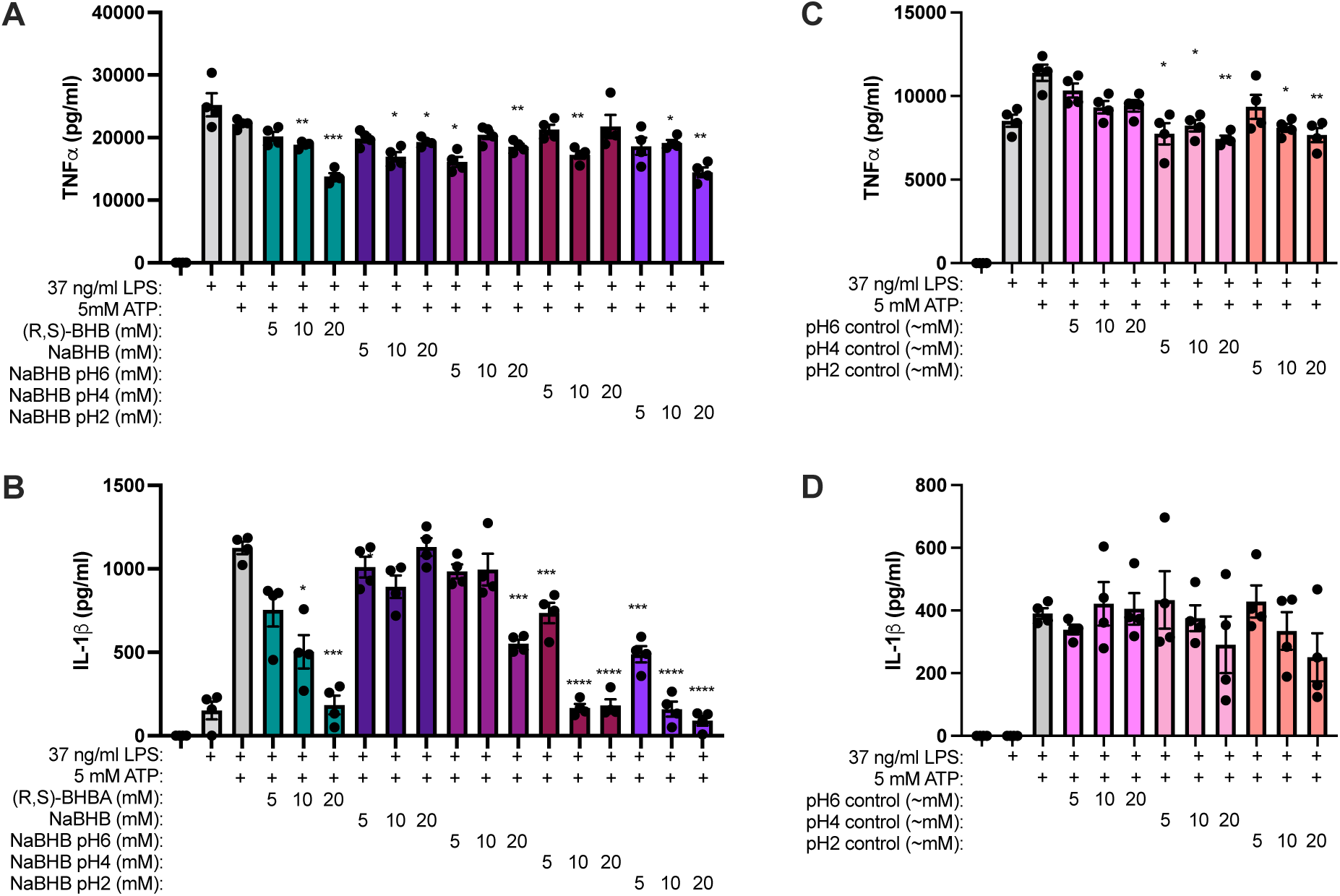
Acidifying the pH of NaBHB confers upon it the capacity to inhibit NLRP3 inflammasome activation, whereas acid alone is insufficient to decrease the abundance of soluble IL-1β. J774.1 macrophages were unstimulated or primed with LPS for 6 hours then stimulated with ATP in the presence of increasing concentrations of (*R,S*)-BHBA, NaBHB prepared at a neutral pH, or NaBHB prepared as 1M stocks with a pH of 6 (NaBHB pH6), 4 (NaBHB pH4), or 2 (NaBHB pH2) for 1 hour, after which soluble TNFα **(A)** and IL-1β **(B)** were measured from the cell culture supernatants. J774.1 macrophages were unstimulated or primed with LPS for 6 hours then stimulated with ATP in the presence of increasing concentrations of HCl solutions prepared in tissue culture media with pH equivalent to those in the 1M acidified NaBHB solutions (pH6 control, pH4 control, and pH2 control, respectively) for 1 hour, after which soluble TNFα **(C)** and IL-1β **(D)** were measured from the cell culture supernatants. N = 4 samples/group. * = p≤0.05, ** = p≤0.01, *** = p≤0.001, **** = p≤0.0001 compared to LPS+ATP.

### Other acidic ketones inhibit NLRP3 inflammasome activation

Since we have reported that several short-chain alcohols share the capacity to inhibit NLRP3 inflammasome activation (50), we examined the capacity of several molecules of different carbon chain lengths with structures and pKa values similar to that of BHBA (**Table 1**) to affect ATP-driven IL-1β secretion from LPS-primed macrophages. Whereas no compounds substantially altered the abundance of TNFα detected (**Figure 3A**), the 4-carbon molecules BHBA (3-hydroxybutyric acid), butyric acid, and 2-hydroxybutyric acid, the 3-carbon molecules 3-hydroxypropanoic acid and 2-hydroxypropanoic acid (lactic acid), the 5-carbon molecule 3-hydroxyvaleric acid, and the 6-carbon molecule 2-hydroxyhexanoic acid all dose-dependently inhibited IL-1β release (**Figure 3B**). In contrast, the amino-modified 4-carbon molecule 4-NH_3_-3-hydroxybutyric acid (or its DMSO vehicle control) was unable to affect the abundance of TNFα or IL-1β (**Figure 3**), implicating particular structural requirements beyond carbon chain length, pKa, and hydroxylation position for the NLRP3 inflammasome-inhibiting effects of these compounds. The requirement of acidification was tested by neutralizing the 1M stocks of these NLRP3-inhibiting compounds with NaOH to pH 7, which did not substantially affect the abundance of TNFα (**Figure 4A**) but did substantially abrogate their ability to decrease IL-1β concentrations (**Figure 4B**) compared to the stock solutions that were not neutralized.

**Figure 3.**
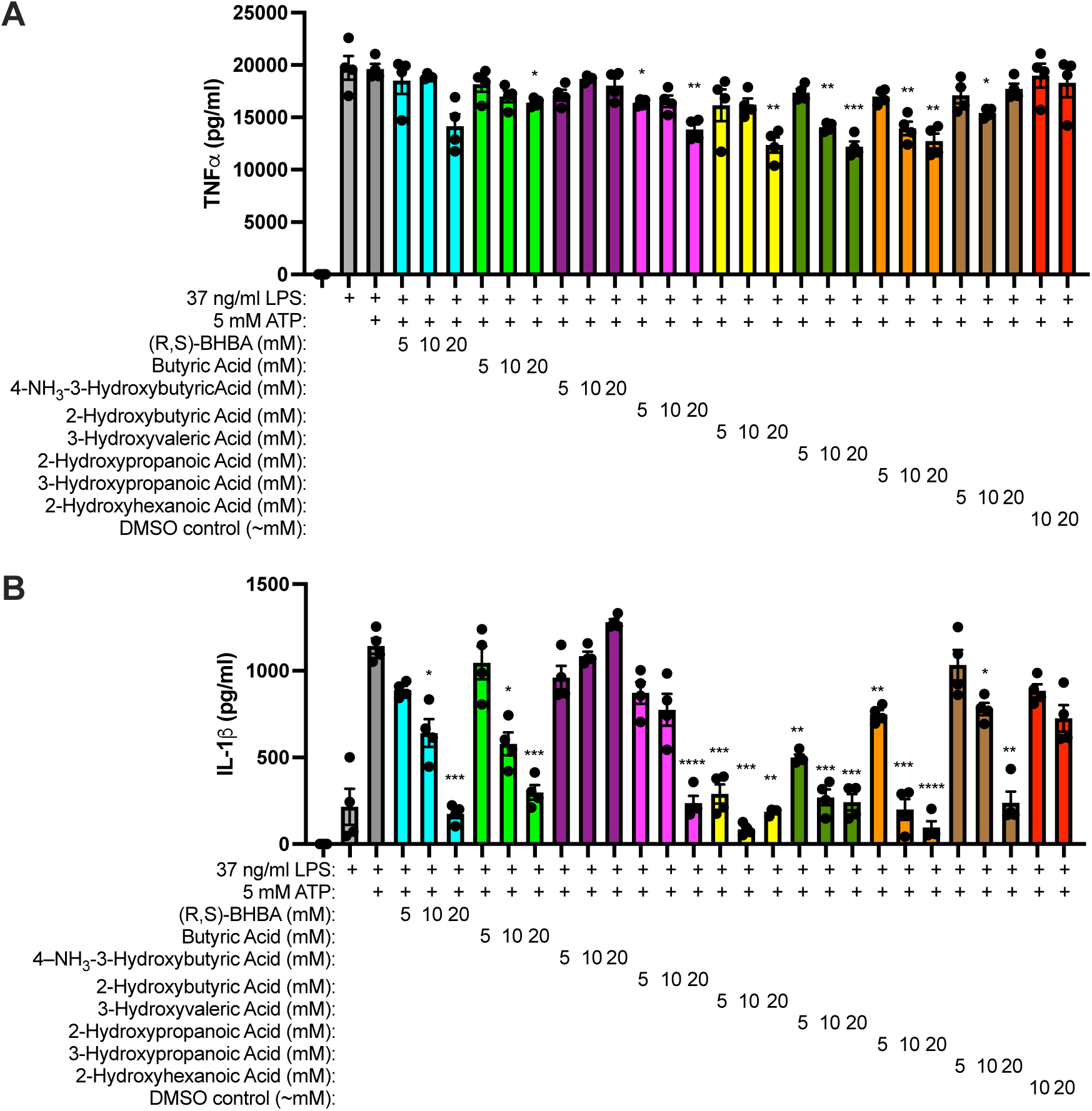
Other acidic ketones inhibit NLRP3 inflammasome activation. J774.1 macrophages were unstimulated or primed with LPS for 6 hours then stimulated with ATP in the presence of increasing concentrations of (*R,S*)-BHBA or several other molecules with ‘BHBA-like’ ketone body structures (or vehicle controls for the concentration of DMSO required to solubilize 4-NH_3_-3-Hydroxybutyric Acid in a 1M stock solution) for 1 hour, after which soluble TNFα **(A)** and IL-1β **(B)** were measured from the cell culture supernatants. N = 4 samples/group. * = p≤0.05, ** = p≤0.01, *** = p≤0.001, **** = p≤0.0001 compared to LPS+ATP.

**Figure 4.**
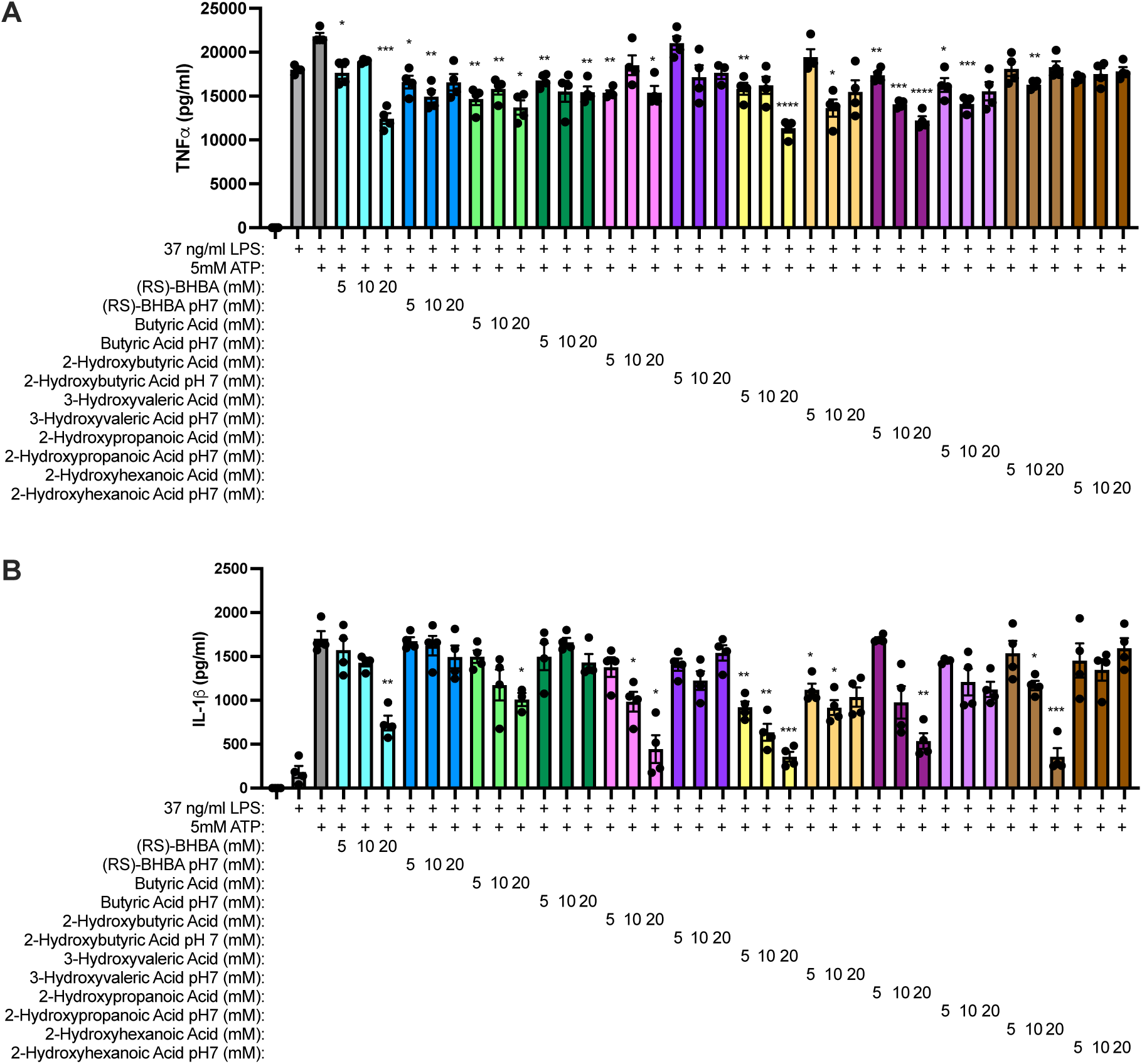
Neutralizing other acidic ketone bodies diminishes their NLRP3 inflammasome-inhibiting effects. J774.1 macrophages were unstimulated or primed with LPS for 6 hours then stimulated with ATP in the presence of increasing concentrations of (*R,S*)-BHBA or several other molecules with ‘BHBA-like’ ketone body structures prepared from 1M stocks that were not or were neutralized to pH7 (using NaOH) for 1 hour, after which soluble TNFα **(A)** and IL-1β **(B)** were measured from the cell culture supernatants. N = 4 samples/group. * = p≤0.05, ** = p≤0.01, *** = p≤0.001, **** = p≤0.0001 compared to LPS+ATP.

**Table 1.**
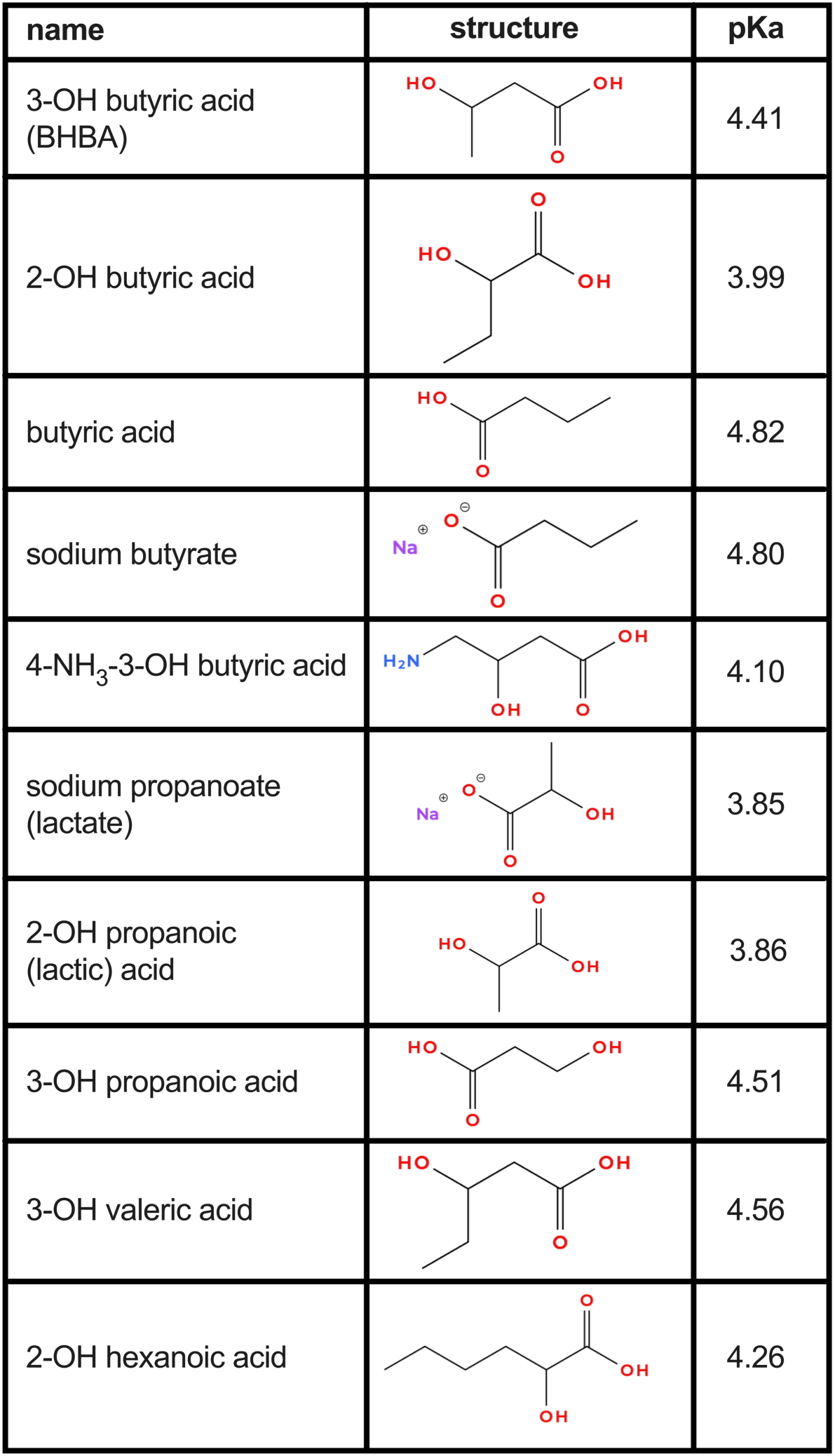
Names, chemical structures, and pKa values of ketone bodies and short-chain carboxylic acids used in these studies. -OH = hydroxy, -NH^3^ = amino. Structures were generated using MolView (https://molview.org).

| name | structure | pKa |
| --- | --- | --- |
| 3-OH butyric acid<br>(BHBA) |  | 4.41 |
| 2-OH butyric acid |  | 3.99 |
| butyric acid |  | 4.82 |
| sodium butyrate |  | 4.80 |
| 4-NH<sub>3</sub>-3-OH butyric acid |  | 4.10 |
| sodium propanoate<br>(lactate) |  | 3.85 |
| 2-OH propanoic<br>(lactic) acid |  | 3.86 |
| 3-OH propanoic acid |  | 4.51 |
| 3-OH valeric acid |  | 4.56 |
| 2-OH hexanoic acid |  | 4.26 |

### Acidic ketone bodies do not kill macrophages and instead inhibit NLRP3 inflammasome activation-induced pyroptotic cell death

Since the TNFa detected in the studies in which LPS-primed J774.1 macrophages are subsequently stimulated for 1 hour with ATP to induce NLRP3 inflammasome activation are produced during the 4-hour LPS priming step (as indicated from the data in Figures 1-4 showing the robust TNFα accumulation when cells are stimulated with LPS alone), the capacity of short-chain carboxylic acids with structures similar to that of BHBA to inhibit IL-1β abundance could be due to their compromising the viability of cells. Consequently, we exposed J774.1 macrophages to several acidic and non-acidic forms of these compounds at 20 mM concentration, DMSO and HCl as ‘vehicle’ negative controls, and digitonin as a positive control, for 24 hours. Whereas digitonin elicited nearly complete cell death when assessed using intracellular trypan blue staining, none of the compounds or vehicles tested compromised cell viability (**Figure 5A**). NLRP3 inflammasome activation in LPS-primed macrophages can induce pyroptosis (1, 2), a caspase-dependent form of cell death during which IL-1β is cleaved and secreted. As acidic forms of BHBA-related compounds attenuate the amount of IL-1β produced, we examined their influence on NLRP3 inflammasome activation-induced cell death. In cultures of LPS-primed J774.1 macrophages, ATP stimulation induced substantial cell death (∼ 60% in an hour), which was attenuated by the acidic compounds BHBA, butyric acid, 2-hydroxybutyric acid, and 3-hydroxypropanoic acid, and 3-hydroxyvaleric acid, but not by the sodium salts of BHB, butyrate, or lactate (**Figure 5B**).

**Figure 5.**
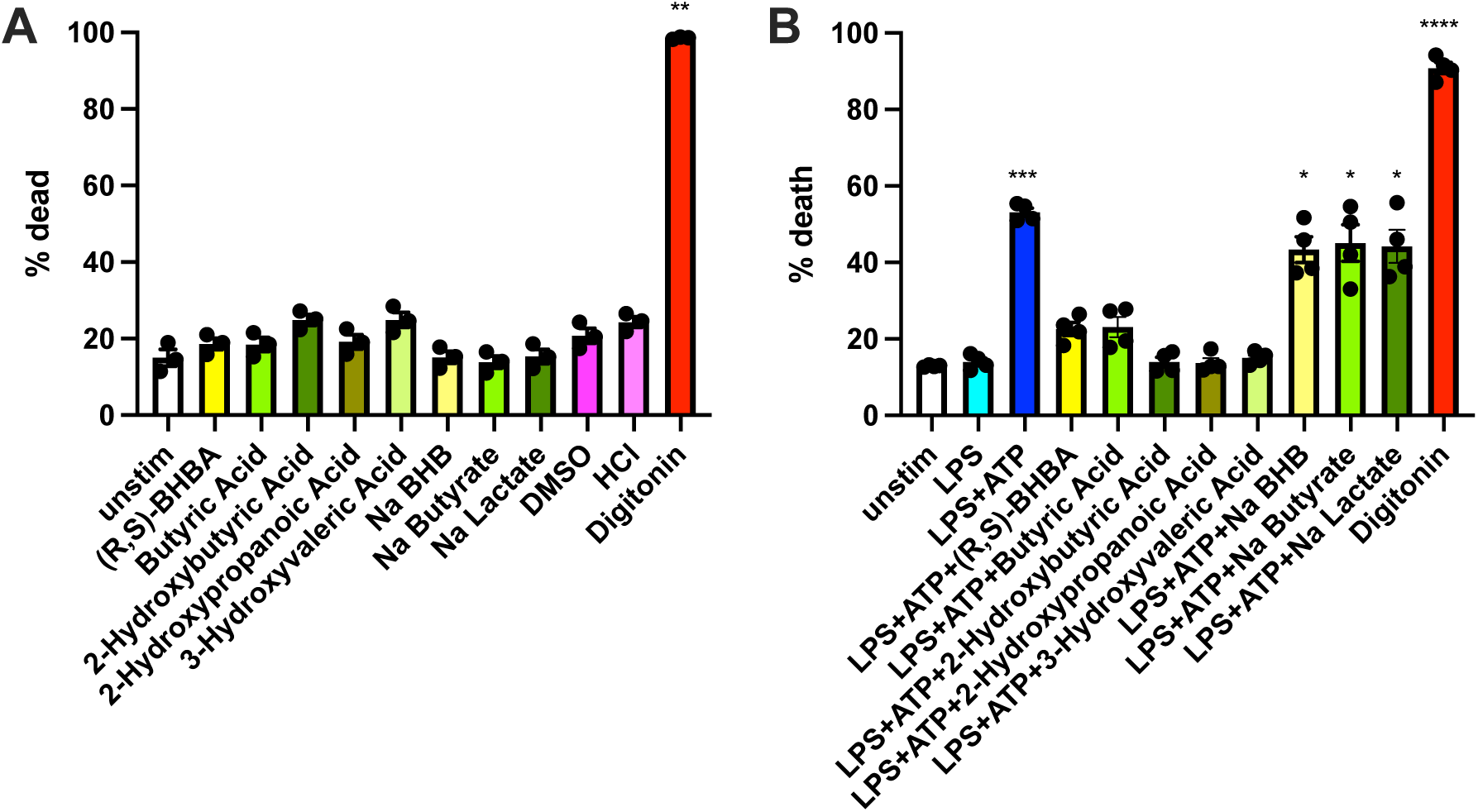
Acidic ketone bodies do not kill macrophages and instead inhibit NLRP3 inflammasome activation-induced pyroptotic cell death. J774.1 macrophages were unstimulated or exposed to 20 mM concentrations of acidic or sodium forms of ‘BHBA-like’ short-chain fatty acids for 24 hours, after which the cells were detached from the cell culture plates by scraping and the percentage of dead cells was assessed using trypan blue staining **(A).** N = 3 samples/group. J774.1 macrophages were unstimulated or primed with LPS for 6 hours then stimulated with ATP in the presence of 20 mM concentrations of acidic or sodium forms of ‘BHBA-like’ ketone body molecules, DMSO and HCl vehicle controls, or 76.5 μM digitonin as an inducer of cell death for 1 hour, after which the percentage of dead cells was assessed by measuring LDH activity from the cell culture supernatants **(B)**. N = 4 samples/group. * = p≤0.05, ** = p≤0.01, *** = p≤0.001, **** = p≤0.0001 compared to unstimulated.

### Acidifying the cellular environment confers NLRP3 inflammasome activity upon NaBHB

We next examined whether modestly acidifying the media of J774.1 cells (in contrast to acidifying the 1M stocks) before NaBHB exposure and inflammasome activation would attenuate IL-1β secretion. Indeeed, while having little effect on TNFα (**Figure 6A**), the addition of a volume of HCl sufficient to influence the pH of the media to the same extent as the pH 2-, pH 4-, and pH 6-adjusted NaBHB used in Figure 2 elicited substantial BHB dose- and acid concentration-dependent inhibition of IL-1β secretion (**Figure 6B**), particularly at the more acidic concentrations. To complement the HCl-mediated acidification experiments, we grew J774.1 macrophages for up to six days in CO2-buffered tissue culture incubators, over which time the pH of the media decreased significantly (**Figure 7A**). Taking advantage of this endogenous metabolic acidification, cells cultured for 6 days were kept in their same endogenously acidic media or washed and provided fresh media. Cells were then primed with LPS and stimulated with ATP in the absence or presence of increasing concentrations of BHBA or NaBHB. Whereas the endogenously acidic condition elicited less cytokine production overall than the fresh media condition the production of TNFα (**Figure 7B**) and particularly IL-1β (Figure 7C) were substantially attenuated by both BHBA and NaBHB. In contrast, only BHBA, not NaBHB, substantially decreased IL-1β secretion from cells stimulated and exposed in fresh media. These results implicate the importance of an acidified cellular environment to confer NLRP3 inflammasome-inhibiting activity to BHB.

**Figure 6.**
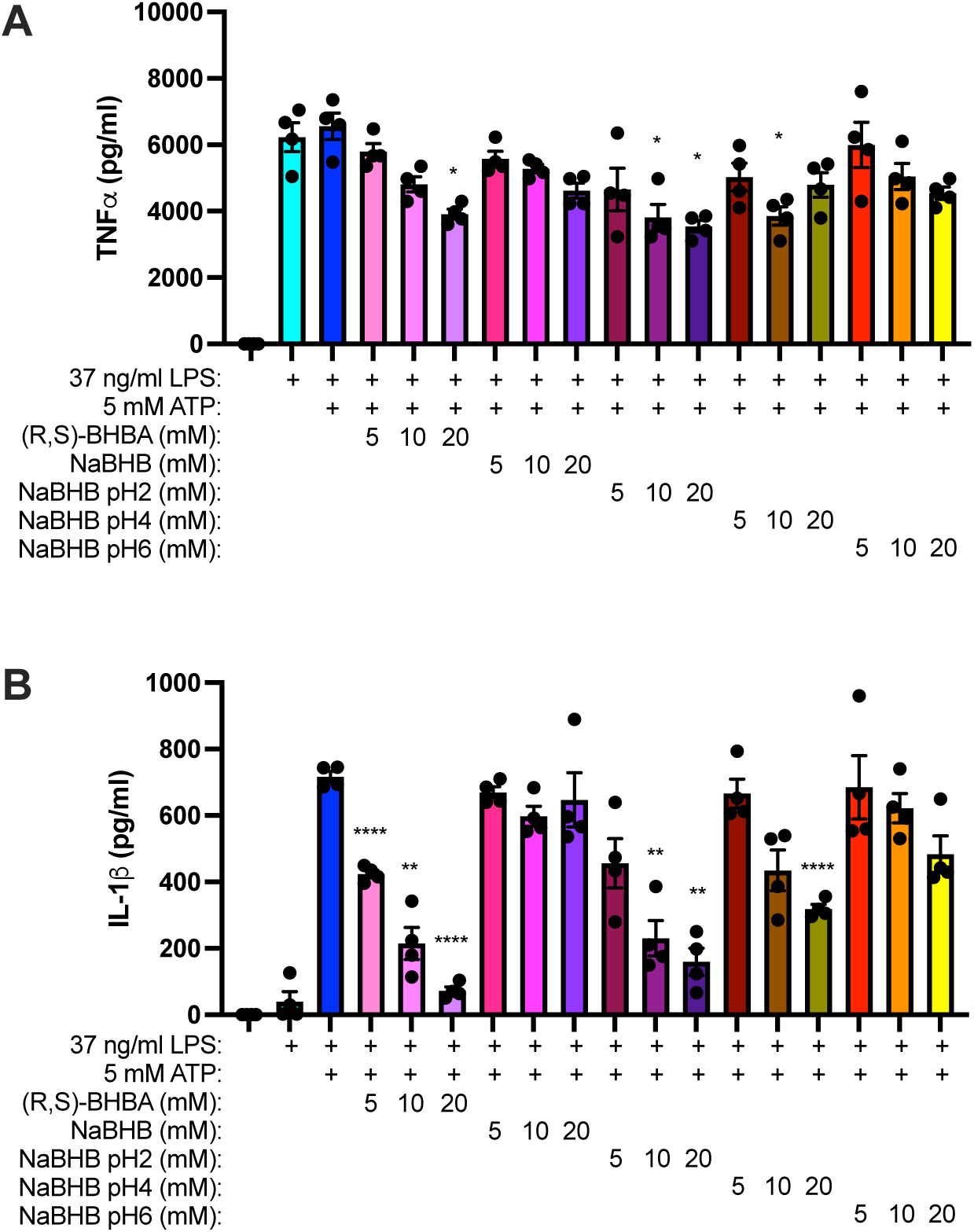
Acidifying the cellular environment confers NLRP3 inflammasome activity upon NaBHB. J774.1 macrophages were unstimulated or primed with LPS for 6 hours then the pH of individual wells was unadjusted or adjusted using 1M HCl to a final pH of 2, 4, or 6, as indicated, and increasing concentrations of (*R,S*)-BHBA or NaBHB were added immediately before cells were stimulated with ATP for 1 hour, after which soluble TNFα **(A)** and IL-1β **(B)** were measured from the cell culture supernatants. N = 4 samples/group. * = p≤0.05, ** = p≤0.01, **** = p≤0.0001 compared to LPS+ATP.

**Figure 7.**
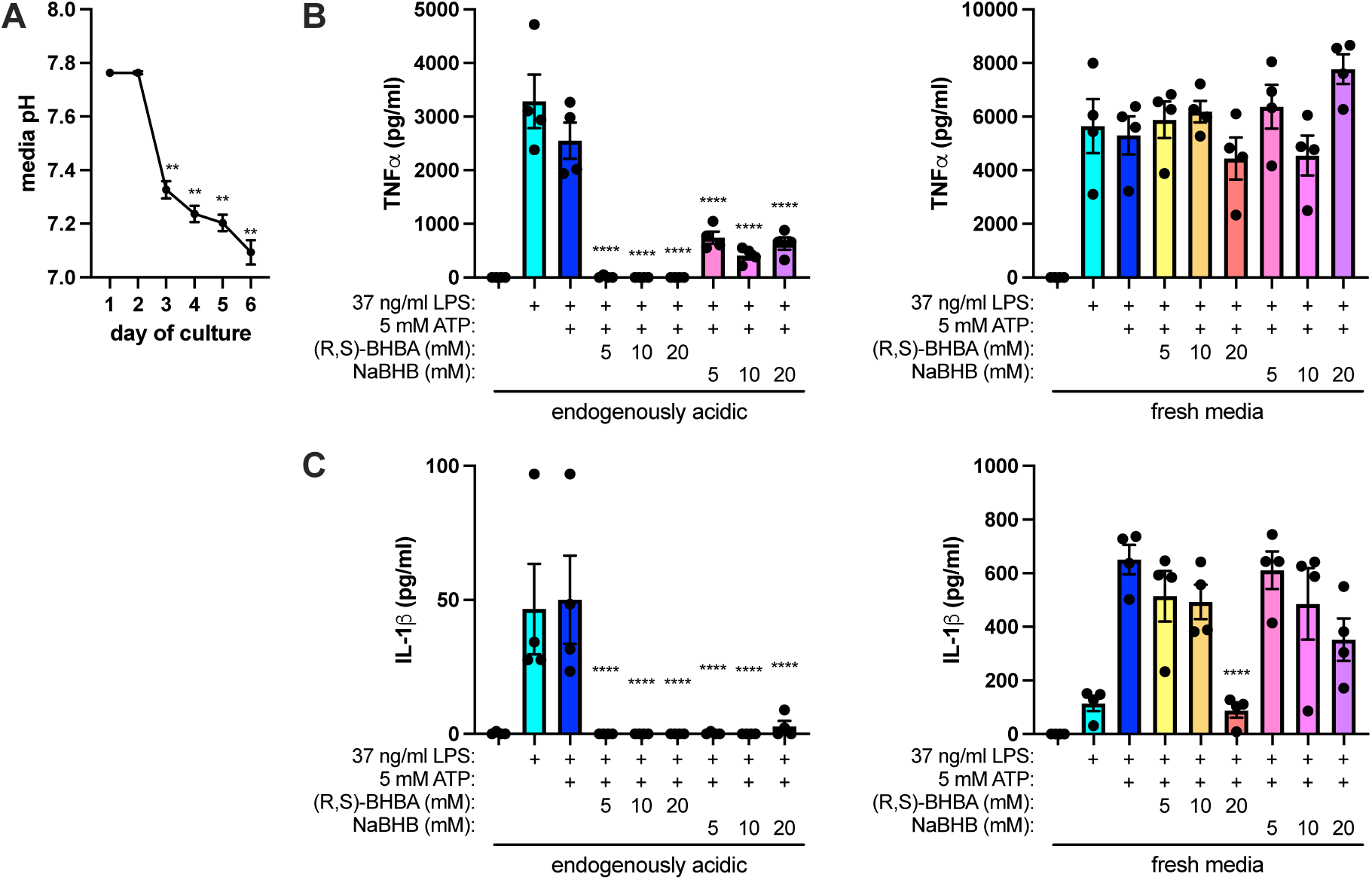
Endogenous metabolic acidification robustly inhibits NLRP3 inflammasome activation. J774.1 macrophages were plated and grown for up to 6 days in a CO_2_-buffered incubator and the pH of the media from individual wells was measured daily **(A)**. J774.1 cells grown for 6 days and maintained in the same media (endogenously acidic) or from which the growth media was aspirated and fresh media was provided, were left unstimulated or primed with LPS for 6 hours then stimulated with ATP in the presence of increasing concentrations of (*R,S*)-BHBA or (*R,S*)-NaBHB for 1 hour, after which soluble TNFα **(B)** and IL-1β **(C)** were measured from the cell culture supernatants. N = 4 samples/group. ** = p≤0.01, *** = p≤0.001, **** = p≤0.0001 compared to day 1 (A) or compared to LPS+ATP in the ‘endogenously acid’ or ‘fresh media’ condition, respectively (B and C).

### Human peripheral blood mononuclear cell (PBMC) NLRP3 activation is inhibited by the acidic forms of β-hydroxybutyrate and similar compounds

To determine whether the effects of ketone bodies observed using mouse macrophages are similar in human monocytes, peripheral blood mononuclear cells obtained from normal donors were unstimulated, primed by exposure for 4 hours to 10 ng/ml LPS, and stimulated with 5 mM ATP for 1 hour to induce NLRP3 activation. In some instances, the LPS-stimulated PBMCs were exposed to BHBA and other acidic compounds, the 1M stocks of which had not been or were neutralized to pH 7 prior to being added to the cells, immediately preceding the addition of ATP. Whereas none of the compounds substantially affected TNF concentrations (**Figure 8A** and **8D**), IL-1β (**Figure 8B** and **8E**) and IL-18 (**Figure 8C** and **8F**) concentrations were dose-dependently inhibited by the acidic but not the neutralized forms of BHBA, butyric acid (BA), 2-hydroxybutyric acid (2-HBA), 2-hydroxypropanoic acid (lactic acid; LA), 3-hydroxyvaleric acid (3-HVA), 2-hydroxyhexanoic acid (2-HHA), and 3-hydroxypropanoic acid (3-HPA). In addition, LPS-primed PBMCs were ATP-stimulated for 1 hour in the presence of increasing concentrations of NaBHB or NaLA prepared from 1M stocks that were unadjusted or were HCl-acidified to pH3. Whereas soluble TNFα concentrations were largely unaffected **(Figure 8G)**, IL-1β and IL-18 concentrations secreted by the PBMCs were substantially inhibited by the acidified NaBHB and NaLA., and the neutral, sodium formulations of BHB (NaBHB) and LA (NaLA) exhibited no IL-1β-or IL-18-inhibiting effects, adjusting the pH of the stocks of these ketone-like compounds to pH 3 using HCl enabled them to dose-dependently decrease the abundance of IL-1β **(Figure 8H)** and IL-18 **(Figure 8I)** in the supernatants of LPS-primed and ATP-stimulated cells. Notably, an equivalent concentration of HCl was unable to decrease the abundance of IL-1β or IL-18.

**Figure 8.**
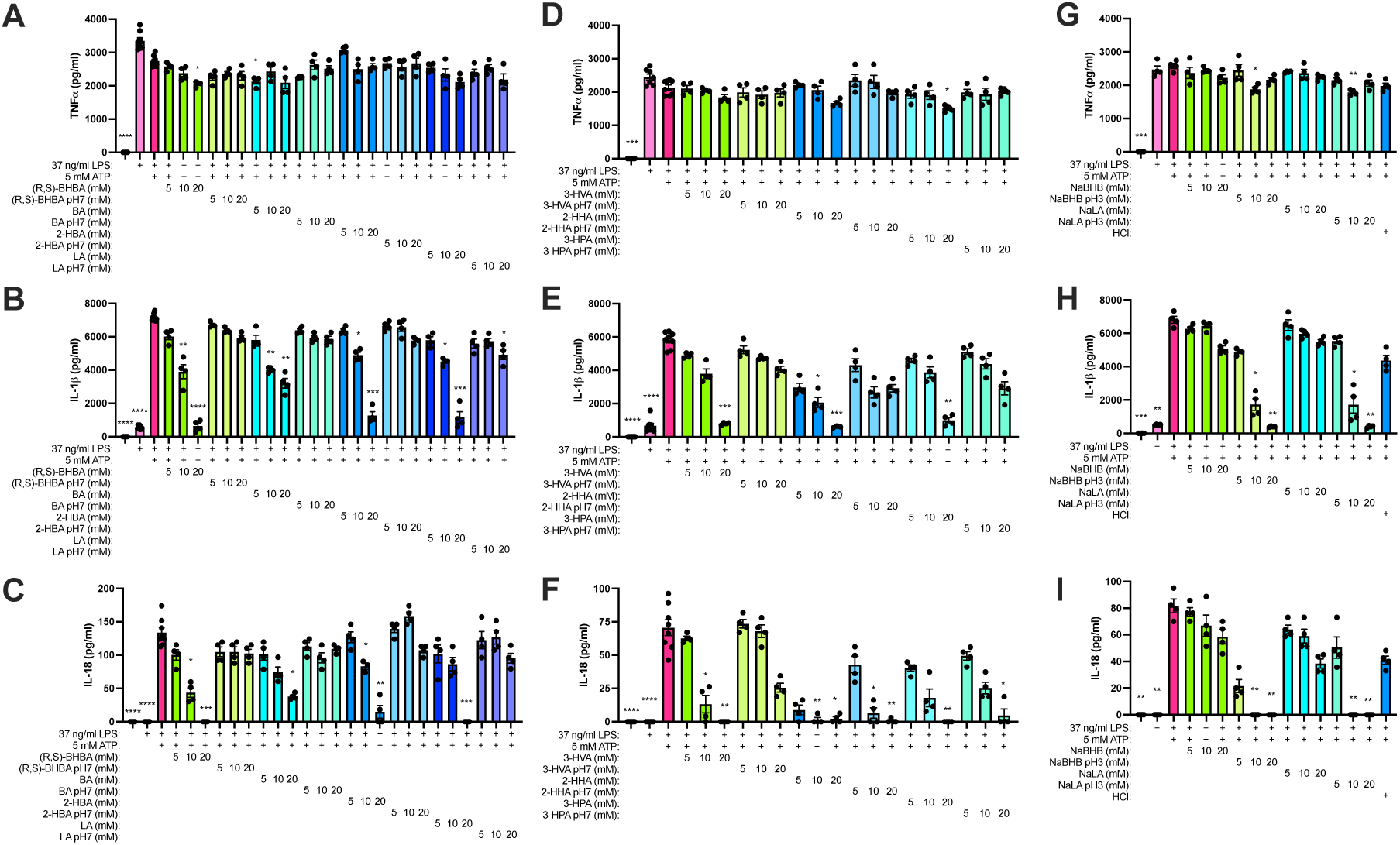
Human peripheral blood mononuclear cell (PBMC) NLRP3 activation is inhibited by the acidic forms of β-hydroxybutyrate and similar compounds. Human peripheral blood mononuclear cells (PBMCs) from normal donors were unstimulated or primed with LPS for 6 hours then stimulated with ATP in the presence of increasing concentrations of (*R,S*)-BHBA or several other molecules with ‘BHBA-like’ ketone body structures prepared from 1M stocks that were not or were neutralized to pH7 (using NaOH) for 1 hour, after which soluble TNFα **(A** and **D)**, IL-1β **(B** and **E)**, and IL-18 **(C** and **F)** were measured from the cell culture supernatants. PBMCs were unstimulated or primed with LPS for 6 hours then stimulated with ATP in the presence of increasing concentrations of NaBHB or NaLA prepared from 1M stocks that were unadjusted or were acidified to pH3 (using HCl) for 1 hour, after which soluble TNFα **(G)**, IL-1β **(H)**, and IL-18 **(I)** were measured from the cell culture supernatants. N = 4 samples/group. * = p≤0.05, ** = p≤0.01, *** = p≤0.001 compared to LPS+ATP.

### GPR41/FFAR3 activation phenocopies and augments the NLRP3 inflammasome-inhibiting effects of BHBA

The metabolic utilization of BHB to generate ATP requires its conversion by β-hydroxybutyrate dehydrogenase (BDH1) to acetoacetate, which is then converted to acetyl-CoA to enter into the Kreb’s cycle and generate energy(52). Since this enzymatic conversion is much poorer using the (*S*)-BHB enantiomer than the (*R*)-BHB enantiomer (52), yet both enantiomers equivalently inhibit NLRP3 inflammasome activation (Figure 1 and (41)), we explored cell surface receptors and transporters implicated to mediate cellular effects of BHB. As the Monocarboxylate Transporter 1 (MCT1) has been reported to facilitate the uptake of BHB (34, 53–55) and lactic acid (56), we evaluated the impact of the MCT1 inhibitor AZD3965 (AZD) in the presence of these compounds on J774.1 macrophages. Cells were unstimulated, exposed to a high concentration (1000 nM) of AZD3965, or primed with LPS for 6 hours then stimulated with ATP in the presence of 20 mM (*R,S*)-BHBA or 2-propanoic (lactic) acid (LA) along with increasing concentrations of AZD3965 for 1 hour. While the abundance of soluble TNFα was unaffected **(Figure 9A)** and the concentrations of secreted IL-1β were substantially inhibited by 20 mM BHBA and LA **(Figure 9B)**, antagonism of MCT1 with increasing concentrations of AZD3965 did not abrogate their IL-1β-inhibiting effects. These results imply that the BHBA-mediated inhibition of IL-1β secretion are independent of its MCT1-mediated uptake.

**Figure 9.**
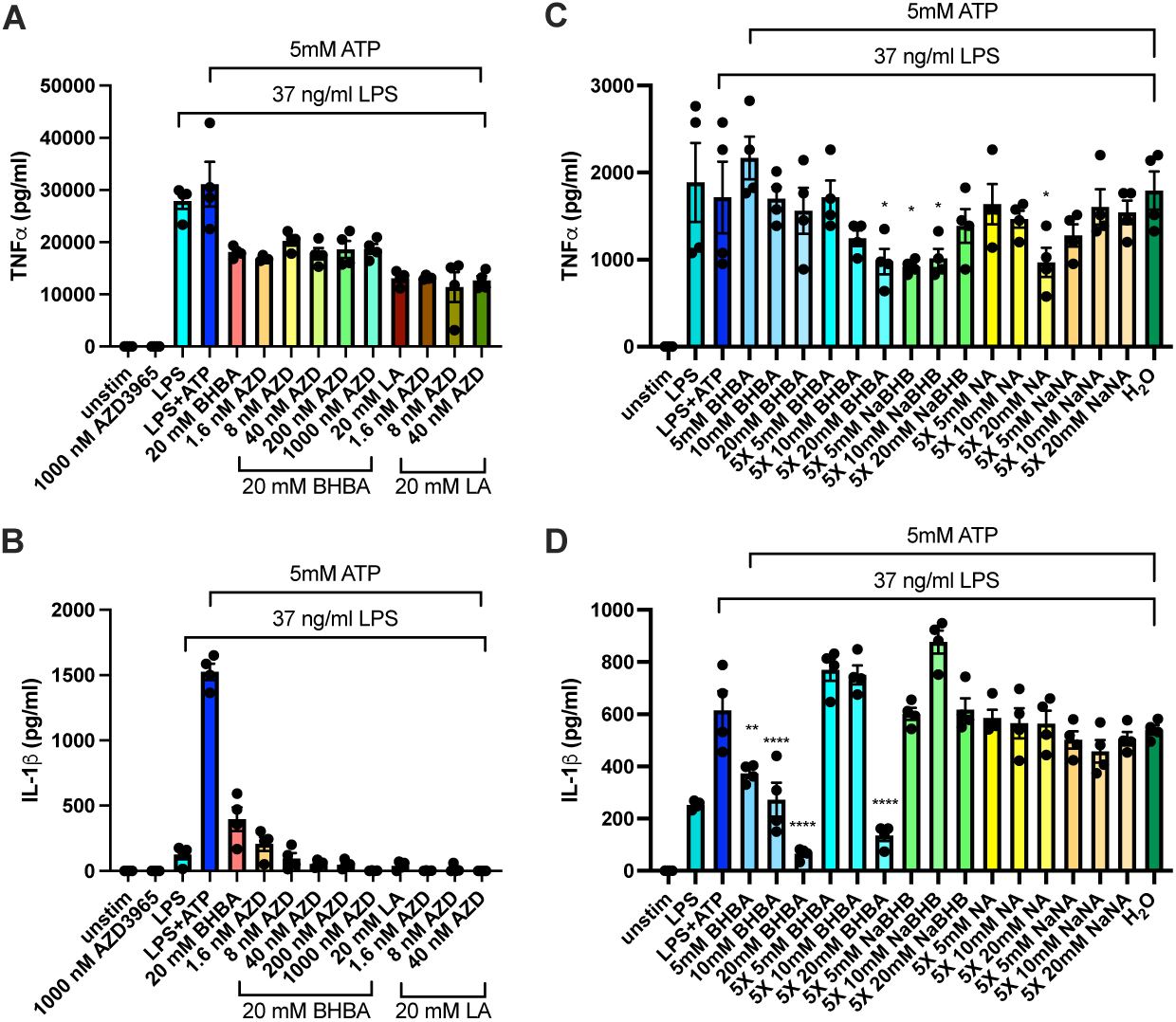
Monocarboxylate Transporter-1 (MCT1) inhibition and GPR109/HCAR2 activation are insufficient to antagonize (block) the NLRP3 inflammasome-inhibiting effects of BHBA. J774.1 macrophages were unstimulated, exposed to a high concentration (1000 nM) of the MCT1 inhibitor AZD3965, or primed with LPS for 6 hours then stimulated with ATP in the presence of 20 mM (*R,S*)-BHBA or 2-propanoic (lactic) acid (LA) in the presence of increasing concentrations of AZD3965 (AZD) for 1 hour, after which soluble TNFα **(A)** and IL-1β **(B)** were measured from the cell culture supernatants. J774.1 macrophages were unstimulated or primed with LPS for 6 hours then stimulated with ATP in the presence of increasing concentrations of BHBA, NaBHB, nicotinic acid (NA), or sodium nicotinate (NaNA), some of which started as 5X more concentrated stocks to account for the additional components added) or with a H_2_O vehicle control, for 1 hour, after which soluble TNFα **(C)** and IL-1β **(D)** were measured from the cell culture supernatants. N = 4 samples/group. * = p≤0.05, ** = p≤0.01, ***= p≤0.001, **** = p≤0.0001 compared to LPS+ATP+20 mM BHBA or 20 mM LA, respectively (A and B) or compared to LPS (C and D).

As BHB has been proposed to elicit effects through activation of the cell surface receptor, hydroxycarboxylic acid receptor 2 (HCAR2) (49, 53, 54, 57–59), we examined whether nicotinic acid, a pharmacological HCAR2 agonist (60), would attenuate NLRP3 inflammasome activation to an extent similar to BHB or LA. J774.1 macrophages were unstimulated or primed with LPS for 6 hours then stimulated for 1 hour with ATP in the presence of increasing concentrations of BHBA, NaBHB, nicotinic acid (NA), or sodium nicotinate (NaNA), some of which started as 5X more concentrated stocks to account for the additional components added, or with a H_2_O vehicle control. Whereas TNFα concentrations were relatively unaffected by the treatments (**Figure 9C**), and BHBA, but not NaBHB, inhibited IL-1β secretion, neither NA nor NaNA inhibited the accumulation of IL-1β in the cell culture supernatants **(Figure 9D)**. These results imply that the BHBA-mediated inhibition of IL-1β secretion is independent of its capacity to activate HCAR2.

Another cell surface receptor reported to mediate the effects of BHB is the Free Fatty Acid 3 (FFAR3) / (GPR41) (53, 54, 58, 59, 61–63). As we have previously reported that BHBA inhibits LPS-induced priming in macrophages (51), J774.1 macrophages were unstimulated or primed with LPS for 6 hours in the absence or presence of increasing concentrations of BHBA without or with the GPR41/FFAR3 activator AR420626. At a BHBA concentration of 10 mM, FFAR3 activation enhanced the inhibition of TNFα secretion. Furthermore, whereas there was little effect on TNFα (**Figure 10B**), when administered to LPS-primed macrophages at the time of ATP stimulation, AR420626 was sufficient to decrease IL-1β secretion to an extent similar to that of 10 mM BHBA (**Figure 10C**). Finally, FFAR3 activation in combination with either 10 mM or 20 mM BHBA elicited an additive effect to inhibit IL-1β secretion from ATP-stimulated cells (**Figure 10C**).

**Figure 10.**
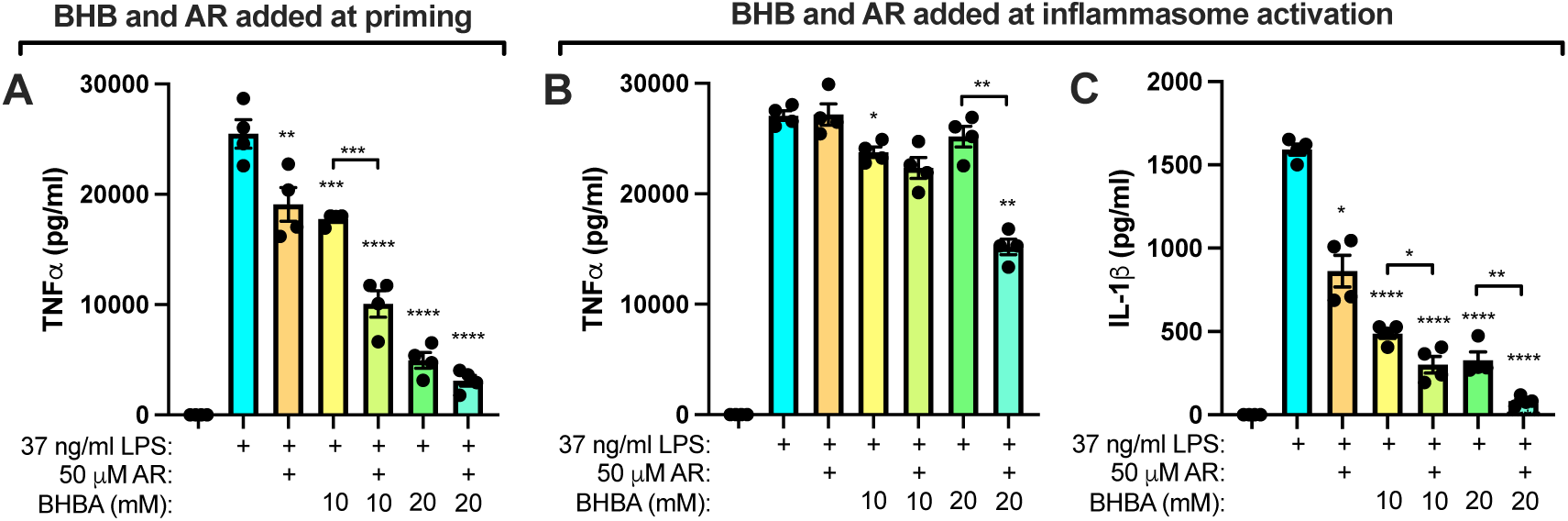
GPR41/FFAR3 activation phenocopies the NLRP3 inflammasome-inhibiting effects of BHBA. J774.1 macrophages were unstimulated or primed with LPS for 6 hours in the absence or presence of increasing concentrations of BHBA in the absence or presence of the GPR41/FFAR3 activator AR420626 (AR), after which soluble TNFα **(A)** was measured from the cell culture supernatants. J774.1 macrophages were unstimulated or primed with LPS for 6 hours then stimulated with ATP in the presence of AR and/or increasing concentrations of BHBA in the absence or presence of the GPR41/FFAR3 agonist AR420626 (AR), after which soluble TNFα **(B)** and IL-1β **(C)** were measured from the cell culture supernatants. N = 4 samples/group. * = p≤0.05, ** = p≤0.01, *** = p≤0.001, **** = p≤0.0001 compared to LPS or as indicated by brackets (A) or compared to LPS+ATP or as indicated by brackets (B and C).

## Discussion

Monocytes in the circulation and macrophages in tissues are components of an integrated system (the immune system) that does not exist in isolation within higher organisms. These and other leukocytes interact with yet other organ systems and are responsive to environmental cues, including those that are nutritional in their origin (64, 65). Reciprocally, metabolism is influenced by leukocyte products, including cytokines (66). Leukocyte activation frequently occurs in the setting of energetic crises, and both the production of pro-inflammatory cytokines and the process of pyroptosis are energetically expensive cellular activities. For example, basal levels of translation are a robust surrogate for cellular energetics (67), and the production of pro-inflammatory cytokines from monocytes and macrophages requires both new protein synthesis and active secretion processes that are incumbent upon cellular energy consumption (68).

Furthermore, cell depletion subsequent to pyroptosis requires the reestablishment of these cell populations through the energetically-demanding processes of hematopoiesis, entry into the circulation, extravasation, migration, activation, etc. In times of energetic crisis, the integrated organ systems engage in cross-talk in which leukocytes restrain their otherwise robust responses, such as NLRP3 inflammasome activation. While there are many mechanisms in place to do so, ketone bodies in an acidic environment may be signals used to restrain these responses. Nutritional insufficiencies are common, and responses are both highly-regulated and diverse throughout evolution (for example, the response to fasting (32)). Systemic and local energetic crises are present in a myriad of acute and chronic human diseases, including food insecurity, obesity, metabolic syndrome, reproduction and nursing, aging, infection, cancer, etc. Understanding the mechanisms and effectors of these responses provides insight into normal biology and opportunities for prevention and therapeutic intervention.

“Therapeutic ketosis” is an alternative and complementary medicine strategy that is both prominent in the popular spotlight and being explored scientifically for its potential to provide benefit in a myriad of disease settings and via several mechanisms (53, 57, 69–71). As reported herein, our studies demonstrate that the ketone body BHB inhibits ATP-induced NLRP3 inflammasome activation from mouse and human monocytes/macrophages, as assessed by the quantity of secreted IL-1β and/or IL-18 and the amount of pyroptotic cell death. Whereas the ability of BHB to inhibit NLRP3 inflammasome activation *in vitro* and *in vivo* has been previously reported (39, 41), our results demonstrate that this capacity is only manifested by the acidic form of BHB (β-hydroxybutyric acid, not sodium β-hydroxybutyrate) or in an acidic environment, whether conferred by the compound itself or the cellular environment in which the cells are exposed to BHB. Furthermore, the acid-dependent inhibition of NLRP3 inflammasome activation is an effect shared by several other tested short-chain carbocyclic acids spanning carbon chain lengths of 3-6 and whether they are β-hydroxylated, α-hydroxylated, or unhydroxylated at these positions.

Endogenous elevations in ketone bodies are oftentimes present in the setting of acidosis, with the most widely recognized conditions being alcoholic and diabetic ketoacidosis (72). In these instances, blood pH decreases while ketone body concentrations elevate substantially. Similar, but more subtle, decreases in pH and increases in ketone concentrations can occur during or soon after exercise (33), fasting (32), low-carbohydrate and low-calorie dieting (73), the taking of weight loss medications (74), and the anorexic sickness response associated with illness that may be mild to severe (75). Sepsis-associated critical illness has been proposed to be a condition in which a failing response to caloric insufficiency is manifest (30, 31), and the augmentation of BHB through the administration of a ketone ester supplement has been reported to prevent muscle weakness in a mouse model of sepsis-induced critical illness (76).

Similarly, impaired ketogenesis has been reported to cause CD4 T cell dysfunction in COVID-19 (77). Ketones beneficially affect CD8^+^ (78) and ψ8 (79) T cells, which are important mechanisms for protection during critical illness caused by respiratory viral infection (80). Perhaps in part through the inhibition of NLRP3 inflammasome activation and pyroptosis, ketone body elevations in the setting of mild acidosis are an endogenous mechanism to limit the inadvertent self-destructive consequences of an inflammatory response. It is compelling that in a randomized, single-blind, placebo-controlled clinical trial, oral administration of BHB alleviated acute respiratory distress syndrome (ARDS) caused by SARS-CoV-2 infection (81). Elevating ketone body concentrations as a therapeutic has been reported to provide benefit in a preclinical model of *Pseudomonas aeruginosa* lung infection (82), and a clinical trial of ketone ester supplementation for *P. aeruginosa*-infected cystic fibrosis patients is currently being conducted (83). Augmented ketone levels have also been implicated to confer protection against several other models of chronic disease, including obese asthma (84), allergic asthma (51, 62), multiple sclerosis (85, 86), hypertension (87), colitis (88), ischemia-reperfusion injury (89), obesity (79, 90), and others.

Ketone bodies have been implicated to influence cells through a number of transporters and cell surface signaling receptors, including HCAR2/GPR109a (49), MCT1 (34, 53–55), and FFAR3/GPR41 (26, 27, 47–49). From our experiments, a mechanism whereby BHB affects macrophages and monocytes *in vitro* involves activation of the cell surface receptor, FFAR3/GPR41. In contrast, the stimulation of HCAR2/GPR109a, the uptake of BHB by MCT1, or the metabolism of BHB into acetate/acetyl-CoA and entry into the Krebs cycle are dispensable for the NLRP3 inflammasome-inhibiting effects of BHB in macrophages. Although it has been reported previously (41), we were unable to measure substantial or consistent inhibition of potassium efflux from cells in which NLRP3 activation-induced IL-1β secretion was decreased by acidic ketone bodies. In addition, in contrast to ethanol (50), BHBA did not decrease the phosphorylation of ASC (not shown). Of the short-chain carboxylic acid compounds evaluated, only 4-NH_3_-3-OH butyric acid, BHBA with an amino group on the 4^th^carbon, was unable to inhibit ATP-induced IL-1β secretion. This compound differs from the others evaluated in that it is a GABA receptor modulator (91). Perhaps GABA receptors, despite also being 7-transmembrane G protein-coupled receptors, do not elicit the same intracellular signals that function to inhibit ATP-induced NLRP3 activation as does FFAR3.

In our studies, we find that several acidic short-chain carboxylic acids with structures similar to BHB and other ketone bodies function similarly to inhibit NLRP3 inflammasome activation and to inhibit pyroptotic cell death, which are not inhibited by acid alone or by the same ketone-like molecules when the acidity is neutralized. Furthermore, acidification of the sodium salts of these molecules is sufficient to confer upon them NLRP3 inflammasome-inhibiting effects. The NLRP3-inhibiting effects appear to be mediated, at least in part, through FFAR3/GPR41, implicating the importance of the acidic component for licensing the anti-inflammatory effects of ketone bodies. Acidosis occurs during endogenous ketone body generation, albeit not from ketone body synthesis per se (92, 93), and so typically accompanies BHB elevations (in contrast to ketogenesis causing acidosis). Perhaps acidification is an important and endogenous component necessary for the anti-inflammatory effects of BHB *in vivo*, an unexplored concept that could be tested by simultaneously administering a buffer such as sodium bicarbonate, as has been done in human studies of ketone body ingestion. Although not as abundant as BHB or acetoacetate, some of the compounds we found able to inhibit IL-1β secretion and pyroptosis are present in normal human plasma (for example, α-hydroxybutyric acid (94)). Perhaps under particular dietary conditions or patterns of nutrient consumption, these other compounds could provide a summative impact in their concentrations and could be able to additively inhibit NLRP3 inflammasome activity. This may make the relatively high concentrations of individual compounds, as were employed in our *in vitro* studies, more relevant to the overall concentrations achievable *in vivo*. Nonetheless, supplementation with exogenous ketones can achieve BHB concentrations in circulation in accordance with the concentrations able to inhibit IL-1β secretion in our *in vitro* studies (95, 96).

There are several additional limitations to our findings. Namely, to ethically provide mechanistic insight, we used *in vitro* studies in a mouse cell line and primary human cells with characteristics of monocytes/macrophages instead of *in vivo* mouse models or human subjects. In addition, our studies used a single NLRP3 agonist (ATP) and measured secretion of inflammasome-dependent cytokines, IL-1β and IL-18. Although we validated the involvement of NLRP3 and gasdermin D in the secretion of these cytokines using the inhibitors MCC950 (97) and disulfiram (98), respectively, we did not examine IL-1β or IL-18 cleavage by western blotting, use a caspase-1 inhibitor, or examine ASC speck formation (50). Interestingly, we found that ionomycin-induced IL-1β release from LPS-stimulated J774.1 macrophages was inhibited by BHBA but not by MCC950 or disulfiram (not shown), implicating effects of acidic ketone-like compounds that are upstream of gasdermin D and NLRP3, and potentially not limited to only conventional inducers of inflammasome activation. Future studies should examine the impacts of pH balancing during “therapeutic ketosis” to examine the impact on NLRP3 activation in models of critical illness to determine whether the protective effects of acidic ketones exhibit reduced efficacy. Furthermore, examining the necessity and/or sufficiency of FFAR3 for the effects of acidic ketone bodies to antagonize NLRP3 activation could be addressed using genetic approaches. BHB is a class-I histone deacetylase (HDAC) inhibitor (26, 27, 99) and its β-hydroxybutyrylation of histone H3 lysines (100, 101) influence gene expression. β-hydroxybutyrylation can also post-translationally modify many other cellular proteins (102, 103). Our efforts to examine β-hydroxybutyrylation in BHB-exposed J774.1 macrophages were inconsistent and inconclusive (104). Finally, the concentrations of ketone-like molecules used in our studies are above those encountered *in vivo*. Nevertheless, these results provide valuable insight into the functions of endogenous ketones and other metabolites under physiological and pathophysiological conditions. Additionally, dietary treatments that increase systemic concentrations of BHB *in vivo*, including ketogenic diet, ketone body precursor feeding, or ketone ester supplementation, while in the setting of mild acidosis, could be used to complement existing supportive therapies for the management of pro-inflammatory conditions.

## Grant Support

This work was funded by US National Institute of Health research grants R01 HL142081, R21 AI186022, and T32 HL076122, research grants from the University of Vermont (UVM) Larner College of Medicine and Department of Medicine, research grants from the UVM Office of Fellowships, Opportunity, and Undergraduate Research, and fellowship grants from the US Department of Education (the CMB-Graduate Assistance in Areas of National Need (CMB-GAANN) Training Grant and the Vermont Space Grant Consortium (VTSGC) Graduate Fellowship Program.

## Disclosures

All authors were supported by grants from the US federal government and University of Vermont, and have declared that no relevant conflicts of interest exist.

## Abbreviations

AcAc: acetoacetate
ANOVA: analysis of variance
ASC: apoptosis-associated speck-like protein containing a CARD
ATP: adenosine triphosphate
BHB: beta-hydroxybutyrate
CARD: caspase activation and recruitment domain
DMEM: Dulbecco’s modified Eagle’s medium
IL-: interleukin-
LDH: lactate dehydrogenase
NLRP3: nucleotide-binding domain, leucine-rich repeat containing proteins (NLR) family pyrin domain containing 3
PBMCs: peripheral blood mononuclear cells
PBS: phosphate-buffered saline
RPMI: Roswell Park Memorial Institute
SCCA: short-chain carboxylic acid
SEM: standard error of the mean
TLC: total lung capacity
TNFα: tumor necrosis factor alpha

## Notes

The authors all declare no conflicts of interest.

### Competing Interest Statement

The authors have declared no competing interest.

